# Neuronal FICD shapes motivated behavior through modulation of dopamine receptor dynamics

**DOI:** 10.64898/2026.10.12.758510

**Authors:** Kristina J. Weaver, Benedict Abdon, Brandon Toth, Alexandra Jack, William Giblin, Christian R. Burgess, Scott D. Pletcher, Matthias C. Truttmann

**Affiliations:** Department of Molecular and Integrative Physiology, University of Michigan, Ann Arbor, MI USA; Michigan Neuroscience Institute, University of Michigan, Ann Arbor, MI USA; Neuroscience Graduate Program, University of Michigan, Ann Arbor, MI USA; Geriatrics Center, University of Michigan, Ann Arbor, MI USA

**Keywords:** dopamine receptor, protein quality control, motivation

## Abstract

Motivated behaviors rely on dopaminergic signaling, which in turn depends on the correct folding and trafficking of receptors through the endoplasmic reticulum (ER). Within the ER, the AMPylase enzyme FICD acts as a molecular rheostat through its modification of the chaperone BiP, dynamically tuning the organelle’s responsiveness to folding perturbations and cellular stress. While the biochemistry of FICD is well-established, its role in complex animal behavior remains poorly understood. Here, we identify FICD as a conserved contributor to the generation of motivated behavior. In *Drosophila*, pan-neuronal knockdown of *dFic* significantly reduces feeding, spontaneous locomotion, and reward-seeking behavior. These alterations occur without disrupting circadian architecture, motor capacity, or homeostatic hunger, indicating a specific loss of motivational drive. Depleting *dFic* within subsets of dopamine receptor-expressing (Dop1R1) neurons recapitulates these behavioral alterations. Transcriptomic profiling reveals that d*Fic* depletion does not induce a canonical unfolded protein response, pointing to a specific post-transcriptional alteration rather than global ER stress. At the molecular level, dysregulated dFic alters the localization and abundance of Dop1R1. Finally, we demonstrate that this regulatory axis is evolutionarily conserved, as loss of FICD similarly diminishes motivated behaviors in C. elegans and mice. Together, these findings implicate FICD in the generation of motivated behavior and link dynamic control of ER proteostasis to dopamine-receptor maturation.

**Significance Statement:** Loss of motivation is a disabling, poorly treated feature of depression, Parkinson’s disease, schizophrenia, addiction, and aging, and it is often attributed to alterations in dopaminergic pathways. Our work points to a distinct and underexplored mechanism of dysfunction: the capacity of neurons to fold and deliver dopamine receptors to their functional sites. By connecting a conserved enzymatic ER switch to motivated behavior across species, this study links cellular protein quality control to neuromodulation. It suggests that behavioral drive depends on how well neurons tune folding capacity to receptor demand. Because G-protein–coupled receptors are the largest class of drug targets, enhancing receptor maturation may offer new therapeutic avenues for motivational disorders.

## Introduction

Animals translate internal physiological states and external cues into goal-directed action, a feat that relies on a chain of biological events extending from the intracellular regulation of neuronal signaling to the outward execution of behavior. Motivated behaviors like feeding, foraging, locomotion, and the pursuit of rewarding stimuli are among the most fundamental of these outputs, and their regulation is under study across species. A central molecule in the generation of motivated behavior is the neuromodulator dopamine, which shapes reward, incentive salience, and the vigor of goal-directed action(1–3). In *Drosophila*, dopaminergic signaling through the D1-like receptor Dop1R1 modulates arousal, locomotor activity, and appetitive behaviors, including sucrose acceptance and feeding(4–6), establishing the fly as a powerful system in which the molecular underpinnings of motivation can be dissected. How the availability and function of dopamine-receptor signaling components are themselves controlled at the molecular level remains incompletely understood.

The output of any neuromodulatory system depends not only on transcription but on the correct folding, modification, and trafficking of its signaling proteins. G-protein–coupled receptors such as Dop1R1 are synthesized and folded in the endoplasmic reticulum (ER), where their maturation and export to the cell surface are gated by ER quality-control machinery(5, 7, 8). The ER chaperone BiP (encoded by *Hsc70-3* in Drosophila; *HSPA5* in mammals) is a central node in this machinery, and its activity is coordinated with the unfolded protein response (UPR), the signaling network that matches ER folding capacity to protein load(9–11). Perturbations in ER protein handling therefore have the potential to change the surface availability of neuromodulator receptors and, in turn, the behaviors they support.

One of the most direct regulators of BiP is the enzyme FICD (also known as HYPE), a Fic-domain protein that reversibly AMPylates BiP(12–14). The covalent transfer of AMP from ATP onto a target residue (AMPylation) is a post-translational modification that FICD establishes to inactivate BiP when folding demand is low; the same enzyme removes the modification to restore BiP activity when demand rises, making FICD a bifunctional rheostat that tunes chaperone activity to ER load(14, 15). The biochemistry of this AMPylation–deAMPylation cycle has been characterized in considerable detail, and Fic-mediated AMPylation has been genetically linked to the UPR and to physiology in several systems, including *Drosophila* vision and adaptation to constant light(16–18). FICD targets beyond BiP remain under investigation, with preliminary evidence suggesting that other chaperones and substrates might be AMPylated. Beyond the ER, FICD influences stress responses and neural processes across metazoans(19–21).

Despite this detailed biochemical understanding, the functions of FICD in directing animal behavior are less well characterized. It is unclear whether an enzyme best understood as a regulator of ER chaperone activity contributes to the generation of complex behavioral outputs, and if so, in which cells it acts and through what molecular targets. The convergence of FICD on ER protein quality control, and the strict dependence of dopamine-receptor signaling on the correct folding and trafficking of receptors through that same ER machinery, raises the possibility that FICD influences behavior by acting on the maturation of neuromodulator receptors.

Here we use neuron-specific knockdown of *Ficd* in *Drosophila* (hereafter, *dFic)* to define the behavioral functions of this AMPylase and the mechanisms through which they arise. We find that pan-neuronal reduction of dFic broadly reduces motivated behaviors, including feeding, spontaneous locomotion, and voluntary ethanol intake. These effects occur without disrupting circadian activity structure, development, motor capacity, or homeostatic drive. We identify specific dopamine-receptor neurons in which *dFic* is required for normal behavior, show through transcriptional profiling that the phenotype occurs without induction of a canonical UPR, and demonstrate that reduction or loss of FICD reduces motivated behavior across worms, flies, and mice. Finally, we link dFic to the trafficking of the dopamine receptor Dop1R1, which is mislocalized when dFic is depleted. Together, these findings identify FICD as a rheostatic enzyme that modulates motivated behavior and connect a well-characterized post-translational modification to dopamine-receptor availability in the nervous system.

## Results

### Pan-neuronal knockdown of *dFic* reduces motivated behaviors in *Drosophila*

To determine whether the AMPylase dFic participates in the generation of behavior, we sought a system in which we could selectively manipulate *dFic* in the brain. *dFic* loss-of-function deletion mutations lead to blindness, and this is rescued by overexpression of *dFic* in glial cells of the visual system(18). Because blindness may confound interpretations of motivational phenotypes, we employed a cell-specific RNAi approach. We depleted *dFic* only in neurons using a pan-neuronal nSyb-GAL4 driver (GMR57C10-GAL4) to express *dFic* RNAi (hereafter nSyb>Fic) and asked whether these animals have altered motivated behaviors compared to genetic controls.

We first assessed food intake, a readout of appetitive drive in flies, using the consumption–excretion (ConEx) assay(22, 23). Over a 24-hour period, nSyb>Fic males consumed significantly less food than controls on both a standard laboratory diet (SY10) and a 20% sucrose-only medium (Figure 1A-B). Females showed the same trend towards reduced consumption, with the strongest deficit on sucrose-only food (Figure 1A-B), indicating that neuronal dFic may modulate behavior in both sexes to varying degrees.

**Figure 1.**
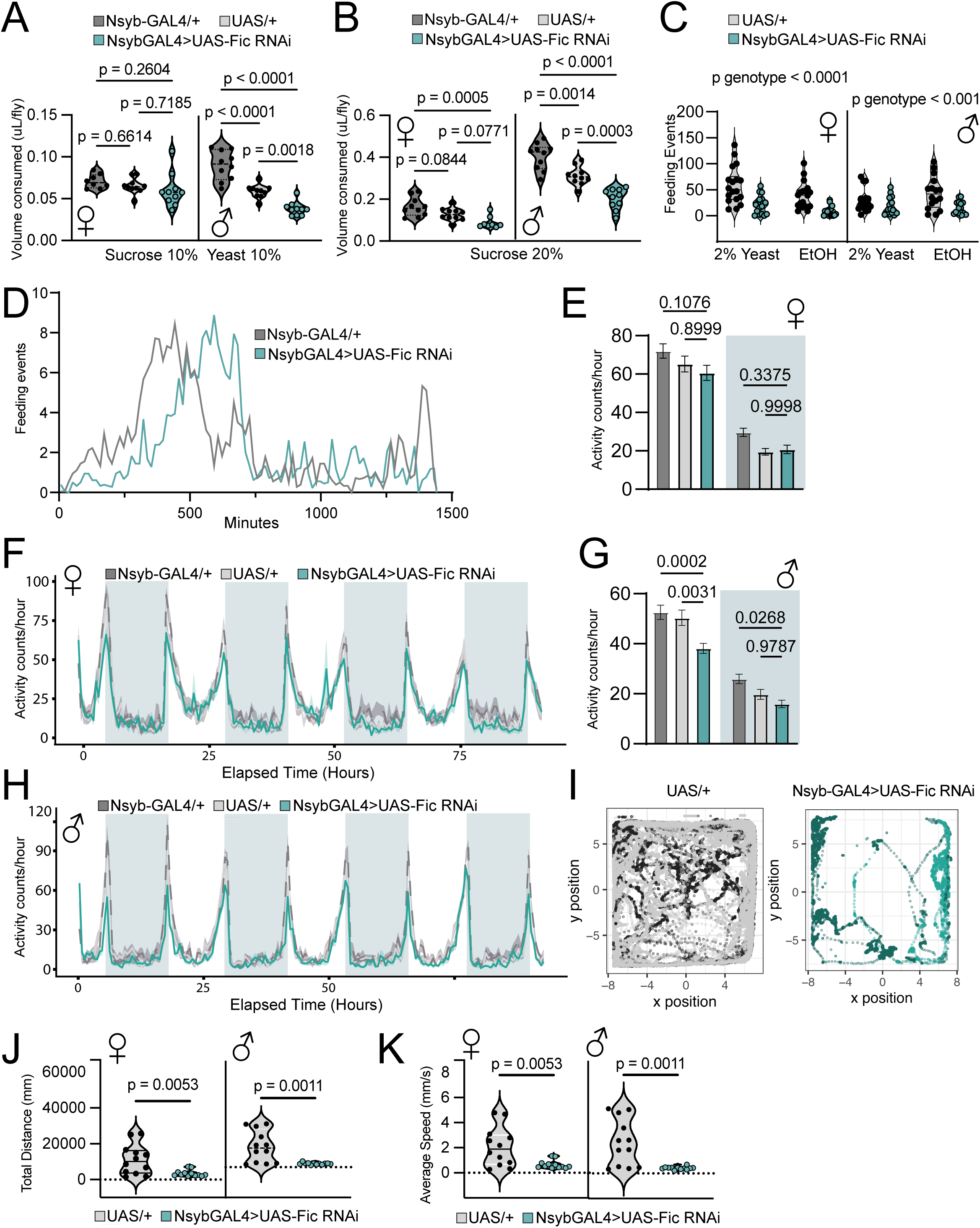
Pan-neuronal knockdown of Fic reduces feeding, locomotion, and reward-seeking without disrupting circadian activity structure. Throughout, genotypes are the pan-neuronal driver control (Nsyb-GAL4/+, dark grey), the RNAi control (UAS-Fic RNAi/+, “UAS/+”, light grey), and the experimental knockdown (nSyb-GAL4>UAS-Fic RNAi, “nSyb>Fic”, teal). **(A)** Food intake of females and males measured by the consumption–excretion (ConEx) assay as feeding volume (µL/fly) on Sucrose 10% and Yeast 10% food [n = 8-10] (one-way ANOVA). **(B)** ConEx feeding volume (µL/fly) on Sucrose 20% food for females and males [n = 8-10] (one-way ANOVA). **(C)** Fly Liquid-Food Interaction Counter (FLIC) interaction events (first 6 hrs quantified) with 2% yeast food versus ethanol-containing food, for females and males (UAS/+ vs nSyb>Fic) [n = 16-18] (two-way ANOVA). **(D)** Feeding-event time course (feeding events vs elapsed minutes) for Nsyb-GAL4/+ and nSyb>Fic over 24 hrs, showing the delayed anticipatory (evening) feeding peak in knockdown flies. **(E)** Female spontaneous locomotion in the Drosophila Activity Monitor (DAM): total activity (counts/hour) separated into light and dark (shaded background) phases [n = 27-29] (two-way ANOVA). **(F)** Female DAM activity time course (counts/hour vs elapsed time, ∼90 h; shaded = dark phase), with preserved morning and evening circadian peaks. **(G)** Male DAM total activity (counts/hour), dark and light phases [n = 27-29] (two-way ANOVA). **(H)** Male DAM activity time course, with preserved circadian peaks. **(I)** Representative single-fly movement traces (x–y position) for a control (UAS/+) and an nSyb>Fic animal, illustrating the reduced spatial coverage of knockdown flies. **(J)** Total distance traveled (mm) from high-resolution video tracking of freely moving females and males (UAS/+ vs nSyb>Fic) [n = 12] (unpaired two-tailed t-test). **(K)** Average speed (mm/s) from the same tracking (unpaired two-tailed t-test). All error bars represent the mean ± SEM.

Feeding behaviors encompass both homeostatic maintenance and the active pursuit of rewarding substances, and we hypothesized that dFic might regulate the drive to interact with a highly rewarding stimulus (24). To test this, we measured voluntary ethanol consumption - a robust, reward-driven appetitive behavior in *Drosophila* - using the Fly Liquid-Food Interaction Counter (FLIC) (24, 25). We presented flies with a choice between two substances to differentiate between behavioral alterations in need-based feeding (homeostatic) versus need-free reward seeking. Flies were given a choice between rewarding ethanol-containing food (15% EtOH + 5% sucrose) or a protein-containing nutritive yeast food (2% yeast extract + 1% sucrose), and we monitored interactions with both for 24 hours. Protein, in the form of yeast, is homeostatically regulated in flies, and thus we reasoned that deficits in need-based feeding would manifest as reductions in feeding only on the yeast food, while deficits in reward seeking would manifest as reductions in both choices(25, 26). nSyb>Fic females and males interacted less overall with both choices compared to controls (Figure 1C), suggesting that dFic is involved in communicating the value of the reward rather than nutritional information.

Consistent with this idea, we also noticed a rightward shift in the feeding peak that occurs in anticipation of their evening meal (Figure 1D). To quantify this shift, we extracted, for each fly, the time of peak interactions and feeding-event rates and the time required to reach half of that peak, and compared these values by genotype using linear mixed-effects models that accounted for the paired ethanol and yeast measurements taken from the same fly. *dFic*-knockdown flies reached their peak interaction rate substantially later than controls (∼552 vs. ∼430 min into the assay) and took roughly twice as long to ramp up to that peak (time-to-half-max: ∼377 vs. ∼185 min; both significant after correction for multiple comparisons). This delay was consistent across both food choices, with no significant genotype-by-food interaction (Supplementary Figure 1 I-J). Notably, the shift in timing was not accompanied by a change in peak amplitude, indicating that the temporal dynamics of feeding engagement and the length of the meal were altered rather than its maximal intensity (Supplementary Figure 1K).

Given that d*Fic* knockdown altered the timing of feeding engagement without changing its peak intensity, we next asked whether this reflects a broader disruption of the circadian machinery that organizes motivated behavior, or whether the timing shift is specific to feeding. To do this, we monitored spontaneous locomotion in the Drosophila Activity Monitor (DAM) assay(27). Both nSyb>Fic females (Figure 1E) and males (Figure 1G) showed decreased overall movement relative to controls, although this did not always reach significance in females. The temporal structure of activity was preserved: the morning and evening activity peaks characteristic of the *Drosophila* circadian program were intact in both females (Figure 1F) and males (Figure 1H), arguing that the deficit is one of activity level rather than a disruption of the circadian clock.

We confirmed and extended these findings with high-resolution video tracking of freely moving flies. Representative traces (Figure 1I) and their quantification (Figure 1J-K) showed that d*Fic* knockdown reduced the total distance traveled in both females and males, and increased micromovements in males (Supplementary Figure 1E). Together, these data show that pan-neuronal reduction of dFic broadly suppresses multiple motivated behaviors - feeding, spontaneous locomotion, and ethanol intake - without disrupting circadian activity architecture.

### Reduced motivated behavior is not explained by developmental, motor, or homeostatic deficits

We next asked whether the behavioral phenotype could be a secondary consequence of impaired development, motor capacity, or sensory or metabolic homeostasis (Supplementary Figure 1). Gross morphological imaging revealed no overt developmental abnormalities in nSyb>fic flies (Supplementary Figure 1A-B), and performance in the negative-geotaxis (climbing) assay was indistinguishable from controls, demonstrating that the animals retain normal motor capability (Supplementary Figure 1C-D). The deficit in feeding and spontaneous movement was therefore not a consequence of an inability to move or to reach food, but perhaps a choice not to.

Several observations indicate that the reduction in feeding reflects diminished motivational drive rather than an inability to sense food or require less of it. When food-deprived, nSyb>Fic flies consumed food at levels equivalent to controls (Supplementary Figure 1F), showing that the machinery for robust feeding is intact and can be recruited under homeostatic demand. nSyb>Fic flies also ate less than controls when tested in the dark (Supplementary Figure 1G), indicating that the phenotype does not arise from an inability to see food. Homeostatic hunger drive, measured by FLIC, was also unchanged (Supplementary Figure 1H)(25, 26). Collectively, these data indicate that neuronal *dFic* knockdown produces a specific reduction in motivated behavior, distinct from developmental, motor, sensory, or homeostatic impairment.

### Pan-neuronal knockdown of *dFic* does not induce a canonical UPR

To identify molecular pathways engaged downstream of neuronal dFic, we performed RNA-sequencing on heads from nSyb>Fic flies and both genetic controls. Principal-component analysis separated nSyb>Fic heads from both controls along PC1 (42% of variance; Supplementary Figure 2), and filtering for transcripts significantly altered in the same direction relative to both nSyb/+ and UAS/+ controls (padj < 0.05, |log2FC| > 1) yielded a high-confidence set of 97 differentially expressed genes (63 down, 34 up; Supplementary Figure 3A).

Given the established role of Fic in AMPylating the ER chaperone BiP and thereby tuning the unfolded protein response (UPR), we examined a panel of canonical UPR components directly (*Ire1*, *Pek*, *Atf6*, *Xbp1*, *crc*, *Hsc70-3*, *Pdi*, *Hrd3*, *sip3*). None reached significance relative to both controls; the only gene trending toward significance was *Hsc70-3*, the *Drosophila* BiP ortholog, which was modestly *downregulated* relative to nSyb/+ alone (log2FC = −0.38, padj = 0.011; not significant vs. UAS/+) (Figure 2A, Supplementary Table 1). Gene Set Enrichment Analysis reinforced this picture: the transcriptome showed a coordinated *upregulation* of oxidative-energy-metabolism programs (aerobic respiration, oxidative phosphorylation, electron transport chain, mitochondrial translation; NES up to ∼2.5) and a broad *downregulation* of neurodevelopmental and morphogenetic programs (NES as low as ∼−2.3), but no induction of the canonical UPR (Figure 2B). Knockdown of neuronal *dFic* is therefore not accompanied by a transcriptional ER-stress program, suggesting that its consequences may be borne at the level of protein handling.

**Figure 2.**
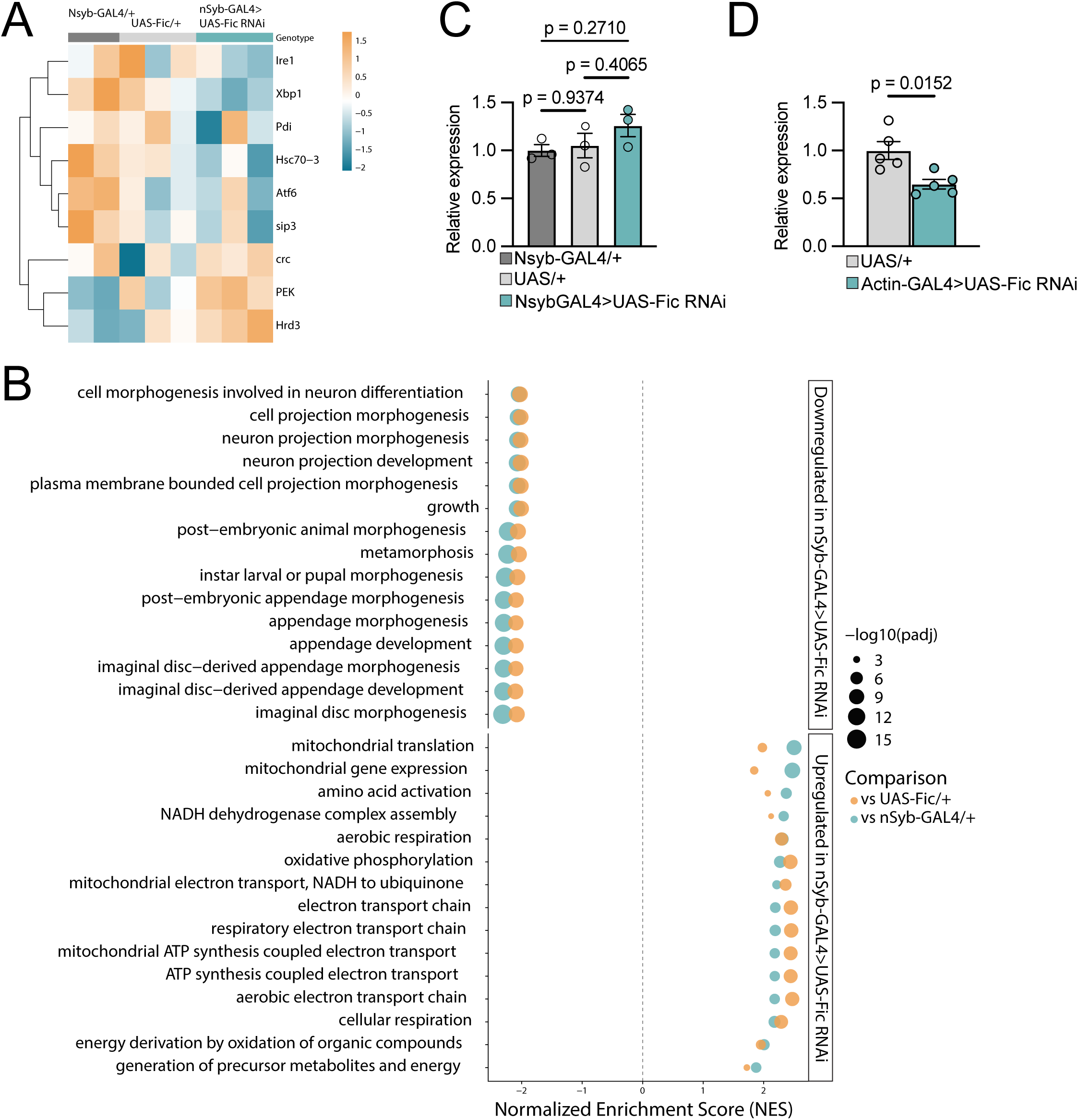
Neuronal Fic knockdown reshapes the head transcriptome toward oxidative metabolism and away from neurodevelopmental programs, without inducing a canonical UPR. **(A)** Heatmap of canonical unfolded protein response (UPR) components (*Ire1, Xbp1, Pdi, Hsc70-3, Atf6, sip3, crc, PEK, Hrd3*; rows, z-scored expression) across the three genotypes (columns). No component is significant relative to both controls. **(B)** Gene Set Enrichment Analysis (GSEA) of GO Biological Process terms, run independently against each control and shown for terms concordant in both (padj < 0.05). Points are colored by comparison (vs. UAS-Fic/+, orange; vs. Nsyb-GAL4/+, teal) and sized by −log10(padj); x-axis is normalized enrichment score (NES). **(C)** qPCR of *dFic* transcript in heads (relative expression) of UAS-Fic/+ versus nSyb-GAL4>UAS-Fic RNAi [n = 3] (one-way ANOVA). **(D)** qPCR of *dFic* transcript in heads of UAS-Fic/+ versus Actin-GAL4>UAS-Fic RNAi [n = 5] (one-way ANOVA). All error bars represent the mean ± SEM. One biological replicate is 10 pooled fly heads.

An unexpected finding of our RNA-sequencing was that d*Fic* transcript was unchanged in nSyb>Fic heads relative to controls, which we confirmed by qPCR (Figure 2C). Bulk-head RNA is dominated by non-neuronal tissue, so this signal is perhaps expected for a neuron-restricted knockdown. Driving the same RNAi ubiquitously with an Actin-GAL4 driver, however, reduced head d*Fic* transcript by ∼40% (Figure 2D), confirming that the RNAi construct is effective when broadly expressed; it has also been previously reported to reduce dFic protein abundance by ∼80% when ubiquitously expressed(18). Together, the absence of a UPR signature and the intact bulk d*Fic* transcript indicate that neuronal dFic reduction may act post-transcriptionally rather than through a transcriptional stress program.

### dFic acts in Dop1R1-receptor neurons to coordinate motivated behavior

We next asked where in the nervous system dFic acts to support motivated behavior. The dependence of appetitive behaviors on dopaminergic signaling led us to test whether dFic is required within dopamine-receptor–expressing neurons to coordinate motivated behaviors. We restricted d*Fic* knockdown to Dop1R1-expressing neurons using a Dop1R1-GAL4 driver (hereafter Dop1R1>Fic, schematic summary of differences from nSyb-GAL4 is depicted in Figure 3A.)(28). This manipulation recapitulated the pan-neuronal feeding phenotype, reducing food consumption in both males and females (Figure 3B). It likewise reproduced the locomotor phenotype: overall movement in the DAM assay was reduced in both males (Figure 3E) and trended downward in females (Figure 3C), while circadian activity peaks remained normal (Figure 3D, 3F), mirroring the pan-neuronal result.

**Figure 3.**
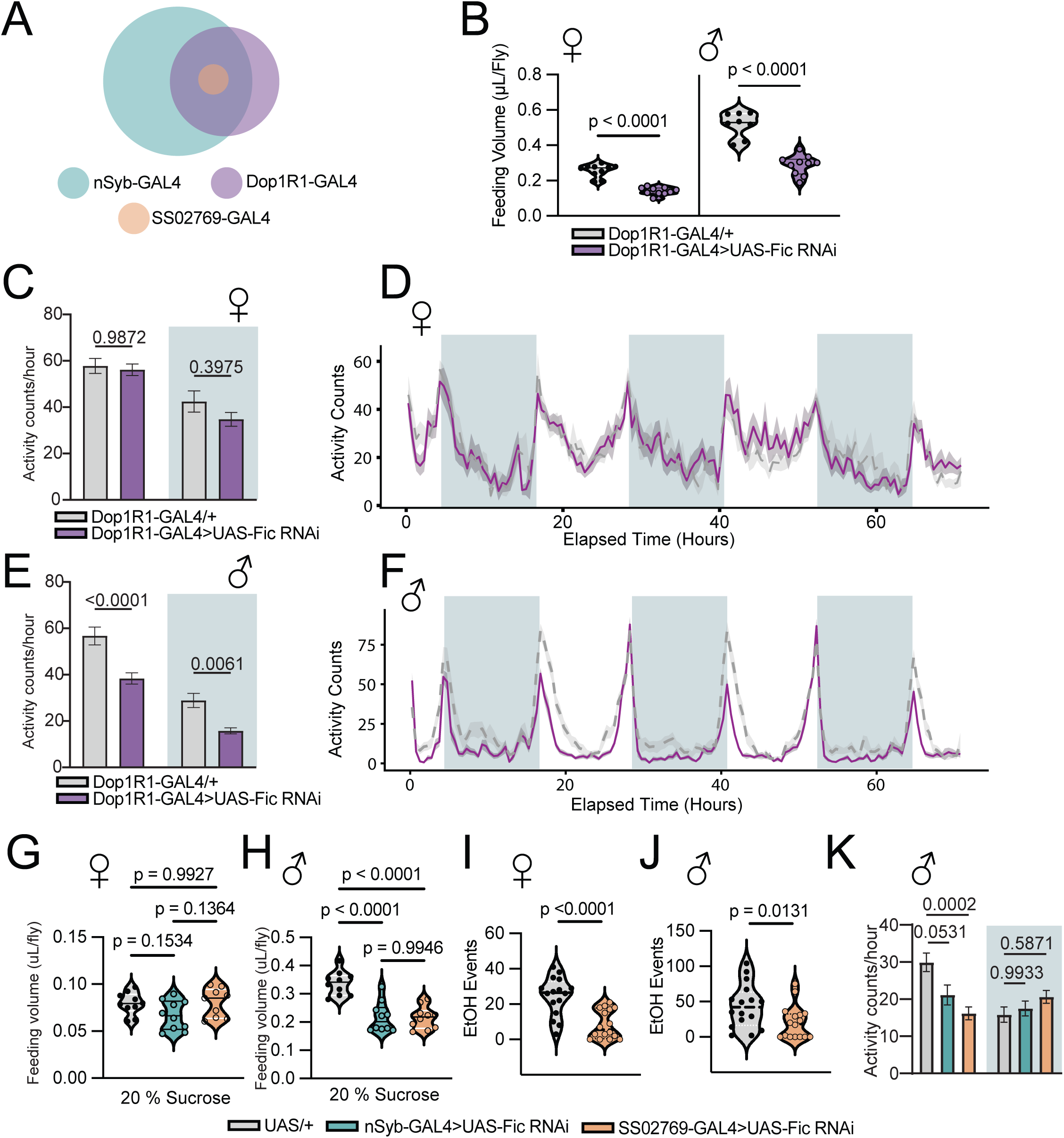
Fic acts within Dopamine receptor-expressing neurons to coordinate motivated behavior. **(A)** Schematic of the adult brain showing the expression domains of the three drivers used, from broad to narrow expression: pan-neuronal nSyb (teal), Dop1R1 (purple), and the SS02769 split-GAL4 (SS02, orange). **(B)** ConEx feeding volume (µL/fly) for females and males: Dop1R1-GAL4/+ versus Dop1R1-GAL4>UAS-Fic RNAi (Dop1R1>Fic) [n = 8-10] (one-way ANOVA). **(C)** Female DAM total activity (counts/hour) separated into dark and light phases for Dop1R1-GAL4/+ versus Dop1R1>Fic [n = 26-29] (two-way ANOVA). **(D)** Female DAM activity time course (counts vs elapsed time; shaded = dark), Dop1R1>Fic (purple) versus Dop1R1-GAL4/+ (grey); circadian peaks preserved. **(E)** Male DAM total activity (counts/hour), dark and light phases [n = 26-29] (two-way ANOVA). **(F)** Male DAM activity time course, circadian peaks preserved. **(G, H)** ConEx feeding volume (µL/fly) on 20% sucrose for the split-GAL4 comparison (UAS/+, nSyb-GAL4>UAS-Fic RNAi, SS02769-GAL4>UAS-Fic RNAi) in females and males [n = 9-10] (one-way ANOVA). **(I, J)** FLIC ethanol interaction events over 6 hrs for the three genotypes in females (I) and males (J) [n = 16] (unpaired two-tailed t-tests). **(K)** Male DAM total activity (counts/hour) for the three genotypes [n = 19-21] (one-way ANOVA). All error bars represent the mean ± SEM.

Dopamine-receptor neurons are found broadly throughout the brain and are functionally heterogeneous, and we wondered whether dFic may be a critical facilitator of behavioral responses within discrete circuit nodes. The R2/R4 ring neurons of the ellipsoid body represent a critical integration center where dopaminergic signaling actively translates external cues into motivated action, having been strongly implicated in mediating cue-induced increases in arousal and appetitive behaviors(4, 5, 29). To test whether dFic acts within this sensorimotor gateway, we used the SS02769 split-GAL4 driver to restrict d*Fic* knockdown exclusively to the R2/R4 subset (Figure 3A, orange)(30). Knockdown of d*Fic* in this population recapitulated the feeding reduction in males (Figure 3H), but not females (Figure 3G), decreased interactions with ethanol in both sexes (Figure 3I-J), and reduced locomotion (Figure 3K). These results identify dopamine-receptor R2/R4 neurons as one example of a cellular locus at which dFic acts to sustain motivated behavior and suggests that there may be sex-specific circuits that regulate diverse aspects of the behaviors.

### Dysregulated dFic activity disrupts Dop1R1 abundance

We next asked how *dFic* knockdown affects proteostasis in the ER. Because dFic’s canonical substrate is an ER chaperone and Dop1R1 is a polytopic G-protein–coupled receptor that matures through the same BiP-dependent ER pathway, we hypothesized that dFic dysregulation impairs the folding or trafficking of the dopamine receptor Dop1R1 itself(31). Western blotting of Dop1R1>Fic fly heads showed a reduction in overall AMPylation (Figure 4A-B). When we quantified Dop1R1, the receptor was increased in both male and female heads relative to controls yet was completely absent in *ΔdFic* null mutants (Figure 4A-B). This increase in Dop1R1>Fic heads was observed in a low as well as a high molecular weight species. We validated the Dop1R1 antibody with a *ΔDop1R1* deletion mutant, in which Dop1R1 protein was also absent, at both the low and high molecular weight. Accumulation of the receptor upon partial knockdown and loss of the mature protein when *dFic* is entirely absent indicate that dFic may be required for the normal maturation and trafficking of Dop1R1 rather than simply for its production.

**Figure 4.**
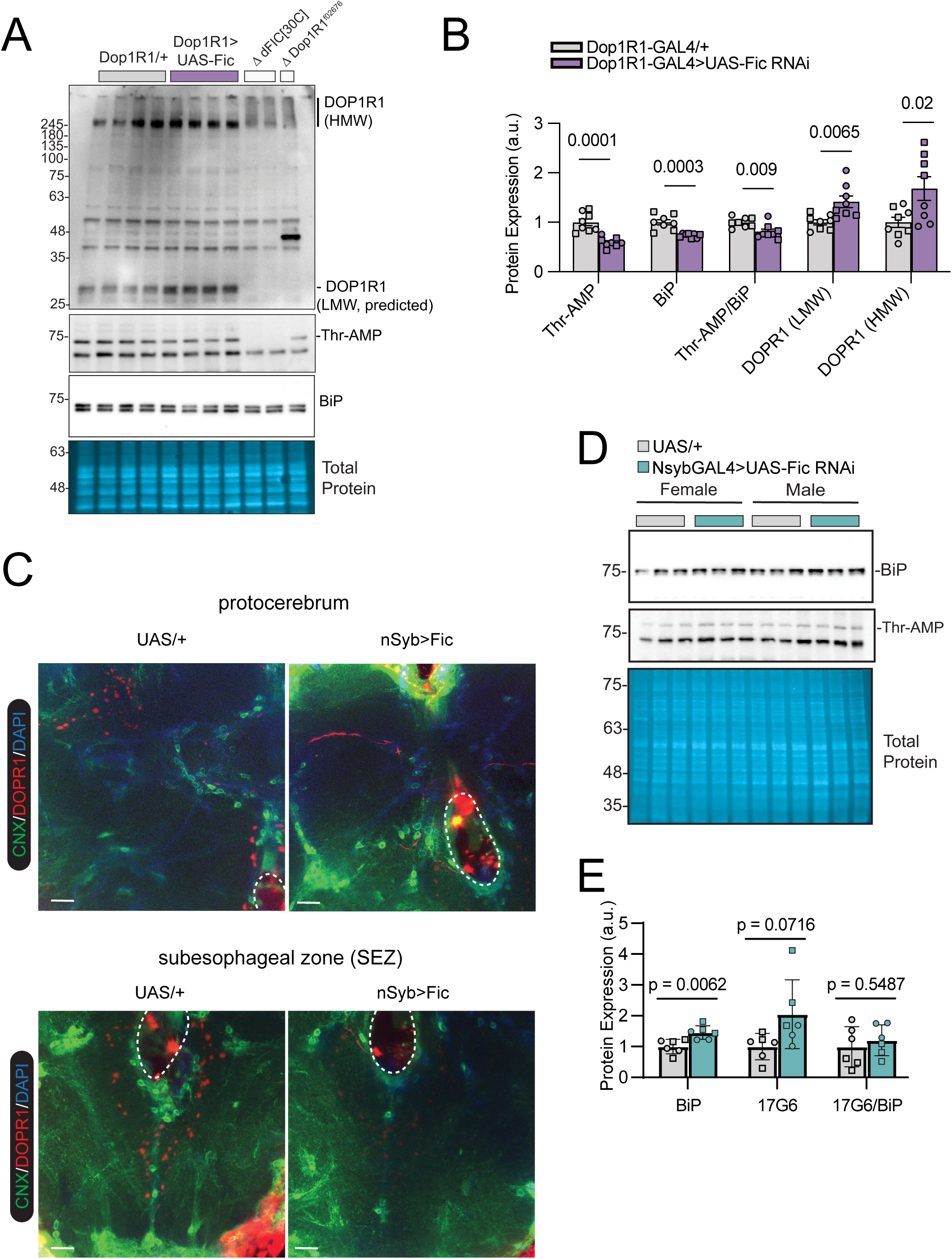
Loss of Fic in neurons reduces and mislocalizes Dop1R1. **(A)** Western blots of male head lysates from Dop1R1-GAL4/+, Dop1R1-GAL4>UAS-Fic RNAi (Dop1R1>Fic), *ΔFic[30C]*, and *ΔDop1R1* knock-in allele (BDSC 84714). Blots (top to bottom): high-molecular-weight species (HMW), Dop1R1 monomer (−DOP1R1, predicted size), BiP, AMPylation (anti-AMPylation antibody 17G6), and total protein (loading control). Molecular-weight markers (kDa) at left. **(B)** Quantification of band intensities (a.u.) across the three genotypes in (A) [n = 8] (one-way ANOVAs). **(C)** Immunofluorescence of UAS-Fic RNAi/+ (control) and nSyb-GAL4>UAS-Fic RNAi brains stained for calnexin (CNX, green), Dop1R1 (DOPR1, red), and DAPI (blue). Protocerebrum (top) and subesophageal zone (SEZ, bottom). Scale bars: 10 µm (n = 3– 5 brains per genotype). **(D-E)** Western blots of head lysates from the pan-neuronal driver (UAS/+ versus nSyb-GAL4>UAS-Fic RNAi) in females and males with quantification of band intensities (a.u.). Squares represent females and circles represent males [n = 6] (one-way ANOVA). All error bars represent the mean ± SEM.

To determine the subcellular basis of this change, we examined Dop1R1 localization by immunofluorescence in nSyb>Fic fly brains. *dFic* knockdown increased total Dop1R1 signal but redistributed it from discrete puncta toward tract-like structures in the subesophageal zone (SEZ) and protocerebrum (Figure 4C, Supplementary Figure 4). This redistribution indicates that, when dFic activity is disrupted, Dop1R1 fails to reach its terminal destination - the hallmark of a receptor trafficking or maturation defect. Together, these findings support a model in which dysregulated dFic-mediated AMPylation interferes with the proper folding and ER export of Dop1R1, linking a conserved post-translational modification to dopamine-receptor availability and, ultimately, to motivated behavior.

One feature of the AMPylation data initially appeared paradoxical. Knockdown with the Dop1R1 driver reduced AMPylation in heads (Figure 4B), whereas knockdown with the pan-neuronal nSyb driver left AMPylation unchanged (Figure 4D-E). Rather than a contradiction, the simplest interpretation is that the two drivers differ in their non-neuronal (e.g. glial) coverage. dFic is expressed in glia as well as neurons(18), so the neuron-restricted nSyb-GAL4 driver leaves this glial pool intact and its knockdown makes little impression on bulk head AMPylation; the Dop1R1-GAL4 driver, which is expressed in ensheathing glia in addition to neurons(32, 33), additionally depletes this reservoir and lowers net AMPylation. Under this model dFic acts as a net AMPylase of BiP in the fly head, and the direction of the bulk change reflects which cells each driver reaches rather than a reversal of Fic’s enzymatic role; a direct test is an important goal for future work. Consistent with a defect in the dynamic control of AMPylation rather than a fixed change in its level, the normal starvation-induced rise in head AMPylation was blunted in Dop1R1>Fic animals, while the bodily response was preserved (Supplementary Figure 5A-D).

### FICD-dependent regulation of motivated behavior is conserved across species

Finally, we asked whether the requirement for FICD in motivated behavior is conserved across species. In *Caenorhabditis elegans*, the loss of *fic-1* is well tolerated in regard to development and lifespan(34). However, we found that *fic-1* null mutants exhibit significantly reduced movement and slower speed in locomotor tracking compared to wild-type controls (Figure 5A-C), mirroring the reduced locomotion seen in flies. Although our fly and worm data point toward a broad reduction in motivated behavior, sucrose preference in mice is a gold-standard behavioral readout for testing reward-seeking directly, making it an important test of whether the phenotype we describe reflects a genuine deficit in motivation rather than a more general effect on movement or intake(35). We found that male, but not female, global FICD knockout mice consumed significantly less sucrose and sucralose compared to wild-type littermates (Figure 5D, Supplementary Figure 6A-C), revealing a more pronounced sex-dependent effect in mice.

**Figure 5.**
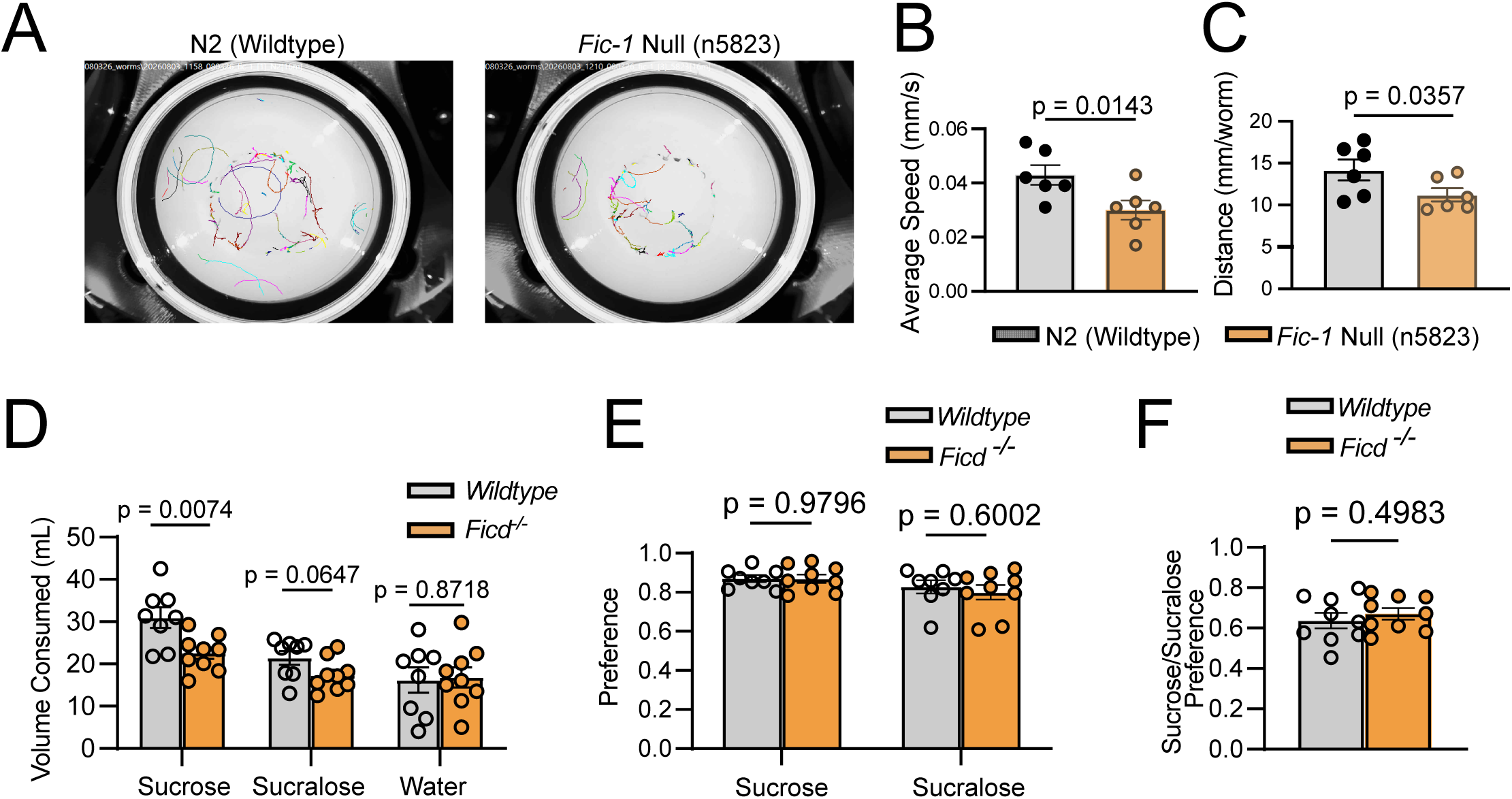
FICD mediates reward-seeking behavior in mice and worms. **(A)** Representative locomotor tracking of C. elegans on plates: N2 (wild-type) and fic-1 null (n5823). **(B)** Average crawling speed (mm/s) for N2 versus fic-1 null [n = 6] (unpaired two-tailed t-test). **(C)** Distance traveled (mm/worm) for N2 versus fic-1 null [n = 6] (unpaired two-tailed t-test). **(D)** Fluid consumption (mL) by wild-type and global Ficd⁻/⁻ (male) mice in a two-bottle test: sucrose, sucralose, and water [n = 8] (unpaired two-tailed t-test). **(E)** Preference (fraction of intake from the tastant bottle) for sucrose and sucralose (unpaired two-tailed t-test). **(F)** Sucrose/sucralose preference ratio (unpaired two-tailed t-test). All error bars represent the mean ± SEM. For C. elegans, one biological replicate is 20-30 worms per genotype.

Control measures confirmed the specificity of this phenotype in males. Water consumption was equivalent between genotypes (Figure 5D, right), and the sucrose preference ratio was likewise indistinguishable between FICD knockout and wild-type animals (Figure 5E), indicating intact relative preference and normal fluid intake. In a two-choice assay, both genotypes and both sexes retained a strong preference for sucrose over the non-caloric sweetener sucralose (Figure 5F, Supplementary Figure 5C), showing that taste-driven choice is also preserved. These findings establish that FICD supports motivated, reward-related behavior in worms, flies, and mice, marking this function as deeply conserved and underscoring sex-specific functions across species.

## Discussion

The biochemistry of FICD-mediated AMPylation has been studied in considerable molecular detail, yet its contribution to the behavior of intact animals has remained undefined. Here we show that FICD is required for the normal expression of motivated behaviors in worms, flies, and mice. Pan-neuronal *dFic* knockdown reduced feeding, spontaneous locomotion, and voluntary ethanol intake in flies while leaving circadian activity structure, development, motor capacity, and homeostatic feeding drive intact. This pattern points to a specific deficit in behavioral motivation rather than a general impairment of health or movement. A loss of behavioral vigor was recapitulated by knockdown of *dFic* in specific dopamine-receptor-expressing populations, and reduction of dFic reduced motivated behavior not only in flies but also in C. elegans and mice. Mechanistically, dFic depletion increased Dop1R1-with the receptor lost entirely in fic-null mutants - and caused Dop1R1 to mislocalize within neurons. Together these findings implicate FICD in the generation of motivated behavior and connect it to the maturation and trafficking of a dopamine receptor.

### A motivational rather than a general behavioral deficit

Several features of the phenotype argue that neuronal dFic supports motivation specifically. The reductions in feeding and locomotion occurred without a change in the temporal organization of circadian activity, and knockdown animals were morphologically normal, climbed normally, and ate as much as controls when challenged with starvation. The preserved response to starvation is particularly informing: the feeding apparatus and the capacity for food intake are intact, but the drive to engage them under baseline conditions is diminished. The specific delay in the timing of the meal peak and the paradoxical increase in micromovements despite reduced total distance travelled further suggest a change in how behavior is organized and initiated rather than in whether it can be executed. This dissociation mirrors the classical distinction between the capacity for a behavior and the motivation to perform it, a distinction that dopaminergic circuits are well known to govern.

### Why loss of an AMPylase would compromise a dopamine receptor: the case for Dop1R1 as a misfolding-prone client

While restricting *dFic* knockdown to Dop1R1 neurons recapitulates the pan-neuronal phenotype and highlights the importance of dFic within dopaminergic circuitry, it remains unclear whether this reflects a unique vulnerability of dopamine receptors or if dFic similarly regulates a broader array of neuronal receptors. Within this specific context, because R2/R4 ring neurons act as a critical gateway for translating external cues into motivated action(36–38), the mislocalization of Dop1R1 may directly limit the capacity of this circuit to initiate cue-induced arousal. When dFic is dysregulated, the resulting trafficking defect effectively prevents Dop1R1 from reaching the sites required to register dopaminergic motivational signals, thereby blunting appetitive drive despite intact homeostatic capability.

Dop1R1 is a class A G-protein–coupled receptor with seven membrane-spanning helices (residues ∼143–450 in *Drosophila* Dop1R1, UniProt P41596). Polytopic membrane proteins of this class are among the most folding-challenged cargoes handled by the secretory pathway: each of the seven transmembrane helices must be correctly inserted, packed, and oriented co-translationally, and the mature receptor must satisfy stringent ER quality-control criteria before export. GPCRs are, as a family, notorious for inefficient folding, with a substantial fraction of newly synthesized receptor degraded rather than exported even under normal conditions, and many disease-associated GPCR mutations act simply by trapping the receptor in the ER(31). Dop1R1 therefore begins as a structurally demanding folding substrate whose surface expression is expected to be sensitive to any alteration in ER folding capacity.

Beyond its transmembrane core, Dop1R1 carries a heavily modified extracellular N-terminus. Sequence analysis identifies an N-terminal ectodomain of ∼142 residues preceding the first transmembrane helix, and this ectodomain contains a cluster of five experimentally supported N-linked glycosylation sites (asparagines N53, N63, N74, N117, and N123; UniProt P41596). N-linked glycosylation in the ER is not merely decorative: the attached high-mannose glycans are read by the calnexin/calreticulin cycle to monitor and assist folding, and glycosylation is frequently required for the ER export and surface delivery of GPCRs(31, 39). A receptor whose folding is chaperone- and glycosylation-dependent is precisely the kind of substrate whose maturation would be compromised when ER chaperone function is perturbed, and whose immature, ER-resident pool would localize abnormally.

### A model linking AMPylation, ER chaperone activity, and receptor trafficking

How does an AMPylase come to control receptor folding? FICD’s best-characterized substrate is the ER Hsp70 chaperone BiP, whose activity FICD reversibly inactivates by AMPylation and restores by deAMPylation, thereby matching active BiP levels to the ER folding load(12–15, 40). BiP, together with the calnexin/calreticulin system, is central to the folding and quality control of nascent secretory and membrane proteins(39). One model is that FICD sets the balance of AMPylated versus active BiP required for efficient folding of demanding clients such as Dop1R1; when FICD activity is dysregulated, this balance is disturbed, Dop1R1 fails to mature, and dopaminergic signaling is altered. An alternative possibility is that Dop1R1 itself may be AMPylated, and this post-translational modification modulates folding and/or transport through direct substrate selection and modification.

An important feature of our transcriptomic data constrains these models: reduction of neuronal dFic did not induce a canonical UPR(9–11). Expression of *Hsc70-3* (*BiP*), *Xbp1*, and *Atf6* was largely unchanged and, if anything, trended downward, and *Gcn2* was unchanged. This argues that the consequences of *dFic* loss are borne at the level of chaperone activity or client handling rather than through a transcriptional ER-stress program, and that the receptor-trafficking defect is not simply a downstream readout of global proteostatic collapse. It is consistent with a model in which AMPylation tunes the functional state of the ER rather than one in which dFic dysregulation triggers a stress response that secondarily damages the cell.

### The direction of BiP AMPylation: interpretation and limitations

A central question our model does not fully resolve is whether the phenotypes reflect an *increase* or a *decrease* in BiP AMPylation in the cells that we manipulated. FICD is a bifunctional enzyme that both AMPylates and deAMPylates BiP, switching between these opposing activities according to ER folding load(14, 15, 40), so reducing its expression does not map cleanly onto a single change in the amount of active BiP. Our two GAL4 drivers changed bulk head measures of AMPylation in opposite directions - unchanged with the neuron-restricted nSyb driver, decreased with the glia-covering Dop1R1 driver - a pattern we speculate is due to differential glial compensation. Even granting this interpretation, whole-head and whole-body immunoblots report the most abundant cell populations and cannot resolve the AMPylation state within the specific neurons that drive the behavior. Nevertheless, a clear functional signature in our data is the loss of *dynamic* regulation rather than a shift in the steady-state level of AMPylation: the normal starvation-induced rise in head AMPylation was blunted in Dop1R1>Fic animals while the bodily response was preserved. This suggests a model wherein folding-demanding clients such as Dop1R1 depend on the capacity to tune BiP activity to changing folding load - a capacity that requires cyclical dFic-mediated AMPylation and deAMPylation and that is disrupted when dFic is perturbed. We therefore interpret our findings as evidence that dysregulated dFic activity interferes with Dop1R1 folding and trafficking.

## Materials and Methods

### Fly stocks and husbandry

Fly stocks were maintained on a standard cornmeal-based larval growth medium and in a controlled environment (21C, 60% humidity) with a 12:12hr light:dark cycle. We used mated female and male flies that were between 7-21 days old for all experiments. The following stocks were used for experiments: *GMR57C10-GAL4* (BDSC 39171), *UAS-Fic RNAi* (v105634)*, Dop1R1-GAL4* (BDSC 84714), *SS02769-GAL4* (BDSC 75997), *Act5C-GAL4 (BDSC 4414), w-;Canton-S, ΔDop1R1^f02676^* (BDSC 605531) and were obtained from Bloomington Stock Center and Vienna Drosophila Resource Center. *ΔFic[30c]* was a gift from Dr. Helmut Kramer. The *Fic* RNAi line used here is a KK library line carrying the hairpin at the 30B landing site only and thus lacks the second 40D insertion that drives non-specific phenotypes in approximately 25% of KK lines (18, 41).

### Mouse rearing conditions

All mouse procedures were conducted at the University of Michigan in accordance with National Institutes of Health guidelines and were approved by the University of Michigan Institutional Animal Care and Use Committee (IACUC), the Animal Care and Use Program, and the Unit for Laboratory Animal Medicine. Wild-type C57BL/6J mice were obtained from The Jackson Laboratory [stock #000664], and whole-body FICD knockout mice were generated as previously described(42). Both lines were maintained as homozygous colonies. Mice were group-housed, up to 5 per cage, under a [12 h:12 h] light/dark cycle, and with ad libitum access to standard chow (Research Diet #5LOD) and water. Male/female mice aged 8-10 months were used for experiments.

### Consumption-excretion feeding assays

Con-Ex experiments were carried out as previously described(23, 43). Blue test food was prepared in removable caps by adding 1% (w/v) FD&C Blue No. 1 to SY10 food or [2% agar + 20% sucrose]. Flies were moved to fresh vials with the removable blue-food caps on the top of the vials and were allowed to feed and excrete for 24 hours. Caps and flies were removed after the 24-hour test period and flies were counted. Excreted dye was collected by vortexing each vial with 3mL of water. Concentration of the dye was determined by absorbance at 630nm and compared to a standard curve of known concentrations.

### Fly Liquid-Food Interaction Counter (FLIC) assays

Flies were tested on the Fly Liquid-Food Interaction Counter (FLIC) system as previously described (24) (Sable Systems International). FLIC *Drosophila* Feeding Monitors (DFMs, Sable Systems International, model DFMV3) were used in the two-choice configuration and each chamber was loaded with food solutions containing a yeast choice ( [2% yeast + 1% sucrose]) or an ethanol choice ([15% ethanol + 5% sucrose]) in 4 mg/l MgCl2. 1.5% Tegosept was added to each food. Flies were briefly anesthetized on ice and aspirated into the DFM chambers, ensuring that each DFM received flies from all treatments. We began recording immediately after loading flies (generally, loading all DFMs requires <10 minutes) and measured FLIC interactions for 24 hours. Total events and temporally binned events were analyzed using custom R code, which is available at https://github.com/PletcherLab/FLIC_R_Code. For each fly and food type, the binned time course was smoothed using locally estimated scatterplot smoothing (LOESS; span = 0.15) to reduce bin-to-bin noise while preserving the overall shape of the feeding response. From each fly’s smoothed trace, two summary features were extracted: Peak Time, the time at which smoothed feeding activity reached its maximum, and Peak Amplitude, the activity level at that maximum. Group differences in each feature were assessed with linear mixed-effects models (lme4 and lmerTest in R), with Genotype, Food Type, and their interaction as fixed effects and a random intercept for individual fly to account for the paired within-animal yeast/ethanol measurements. Fixed effects were evaluated with Type III F-tests, and p-values were corrected for multiple comparisons across features using the Benjamini-Hochberg procedure. Animals that did not participate (i.e. returned zero values), whose DFM returned an unstable baseline signal, or who produced extreme outliers (criteria for outliers were predetermined as exceeding twice the mean of the population)(25) were excluded from analysis.

### Activity Assay

Activity recordings and data processing were performed using the *Drosophila* Activity Monitor System (TriKinetics). Flies were individually tested in 5 mm x 65 mm polycarbonate tubes with SY10 food at one end of the testing tube. The first 12 hours of data was removed from the final analysis in order to allow for acclimation to experimental housing conditions. Total activity counts were calculated for each fly by summing all activity counts recorded during the light and dark cycle, respectively. Experiments were performed at 25°C and 60% humidity under a 12-hour light:12-hour dark cycle.

### Real-Time Video Tracking

Real-time video tracking was conducted using custom equipment and software developed by the Pletcher Lab (DTrack). Pairs of flies of the same genotype were placed into square behavioral arenas measuring approximately 1.5 inches x 1.5 inches. The floor of the arenas were illuminated from below with infrared LEDs to allow for video tracking from above. DTrack locates and tracks individual flies and custom software and code extracts movement data from these traces, which is available at: https://github.com/PletcherLab/ARENA_R_Code.

### Negative Geotaxis

Negative Geotaxis was assayed using custom equipment and software developed by the Pletcher Lab (D-Drop Deluxe and DTrack). Flies were transferred into individual lanes of custom behavioral chambers and then habituated for 10 minutes. Chambers were loaded into the D-Drop apparatus – which drops the chamber containing the flies and then records video of flies climbing up their lanes. Each chamber was dropped 3 consecutive times and recorded for 30 seconds after each drop, and metrics were averaged over 3 trials.

### RNA Sequencing

RNA sequencing was performed on nine biological replicates spanning three genotypes (nSyb-GAL4/+, n = 3; UAS-Fic/+, n = 3; nSyb-GAL4>UAS-Fic, n = 3). Each biological replicate contains 10 heads. RNA was extracted with New England Biolabs Monarch Total RNA Extraction and Isolation kit following manufacturer’s recommendations. Libraries were generated by Plasmidsaurus.

### RNA Sequencing Analysis

Quality of the fastq files was assessed using FastQC v0.12.1. Reads were then quality filtered using fastp v0.24.0 with poly-X tail trimming, 3’ quality-based tail trimming, a minimum Phred quality score of 15, and a minimum length requirement of 50 bp. Quality-filtered reads were aligned to the reference genome using STAR aligner v2.7.11 with non-canonical splice junction removal and output of unmapped reads, followed by coordinate sorting using samtools v1.22.1. PCR and optical duplicates were removed using UMI-based deduplication with UMIcollapse v1.1.0. Alignment quality metrics, strand specificity, and read distribution across genomic features were assessed using RSeQC v5.0.4 and Qualimap v2.3, with results aggregated into a comprehensive quality control report using MultiQC v1.32. Gene-level expression quantification was performed using featureCounts (subread package v2.1.1) with strand-specific counting, multi-mapping read fractional assignment, exons and three prime UTR as the feature identifiers, and grouped by gene_id. Final gene counts were annotated with gene biotype and other metadata extracted from the reference GTF file.

Prior to differential expression analysis, sample quality was assessed using three independent, quantitative diagnostics: principal component analysis (PCA) of variance-stabilizing-transformed (VST) counts, hierarchical clustering of pairwise sample-to-sample Euclidean distances, and per-sample Cook’s distance distributions computed by DESeq2 during model fitting. One sample (nSyb replicate 3) was identified as a clear outlier by all three measures. On PCA, this sample separated from all eight remaining samples along PC1 (58% of total variance), at a distance from the rest of the dataset far exceeding the spread among any other samples regardless of genotype. Consistent with this, pairwise sample-distance analysis showed this sample’s distance to every other sample (∼35–40) was substantially elevated relative to all other pairwise distances in the dataset (∼5–20), indicating it was globally dissimilar to all other replicates rather than differing along a biologically meaningful axis. This was corroborated by Cook’s distance: applying a predefined threshold (mean per-sample median Cook’s distance + 2× interquartile range across all nine samples), this sample had 725 genes with Cook’s distance exceeding 1 — nearly twice as many as any other sample (range across the remaining eight samples: 19–404) — indicating disproportionate influence on model fit across a large fraction of the transcriptome. This sample’s DESeq2 size factor (0.90) was unremarkable relative to its genotype group, indicating the discrepancy reflected an aberrant expression profile rather than a sequencing-depth artifact. One additional sample (Nsyb replicate 1) exceeded the same Cook’s distance threshold (404 genes with Cook’s distance > 1) but was not flagged as an outlier by PCA or sample-distance clustering, and its Cook’s distance elevation was substantially smaller than that of the excluded sample; in the absence of corroborating evidence from independent measures, it was retained in all downstream analyses.

Based on this multi-method assessment, the outlier replicate was excluded, and all subsequent analyses (differential expression, PCA, and gene set enrichment analysis) were performed on the remaining eight samples (nSyb/+, n = 2; Fic/+, n = 3; nSyb>Fic, n = 3).

#### Differential expression overlap analysis

To identify transcriptional changes attributable specifically to neuronal *dFic* knockdown rather than to genetic background effects of either parental control line, differentially expressed genes (DEGs) were independently identified for two comparisons using DESeq2: nSyb>Fic vs. nSyb-GAL4/+ and nSyb>Fic vs. UAS-*Fic*-RNAi/+. Genes were considered significant within each comparison at padj < 0.05 (Wald test, Benjamini-Hochberg corrected) and |log2 fold change| > 1. The overlap between the two significant gene sets was visualized as a Venn diagram. To arrive at a conservative set of genes reflecting active neuronal *dFic* knockdown independent of driver- or transgene-specific effects, the intersection of the two DEG sets was further filtered to retain only genes changing in the same direction (sign of log2 fold change) in both comparisons, yielding a final concordant DEG set of 97 genes (63 downregulated, 34 upregulated) used in all subsequent analyses and figures. For visualization, differentially expressed genes for each comparison were plotted as volcano plots (log2 fold change vs. −log₁₀(padj)), with genes colored by significance and direction using the same padj < 0.05, |log2FC| > 1 thresholds. FlyBase gene identifiers were mapped to gene symbols using the *org.Dm.eg.db* annotation package for labeling the top genes by adjusted p-value in each direction; genes without a corresponding symbol were labeled by their FlyBase ID.

### Quantitative Real-Time PCR

RNA extractions were performed as described above. Complementary DNA (cDNA) was prepared from 1 ug of total RNA using High-Capacity cDNA Reverse Transcription Kit (Thermo Fisher Scientific) and the resulting cDNA was diluted 1:50 before use. No-RT reactions were prepared and run as controls to ensure no genomic DNA contamination. Quantitative polymerase chain reactions (qPCRs) contained 1X PowerUp SYBR Green PCR Master Mix (Applied Biosystems), 500 nM appropriate primers, and 5 ul 1:50 cDNA for a total reaction volume of 10 ul. Relative expression was determined using the comparative CT approach. Primers used in this study are:

*dFic Forward:* CAGAACAACCGAGTCCACCT
*dFic Reverse:* CGCAGTTTCCAGACCAGATT
*Rpl32 Forward:* cgg atc gat atg cta agc tgt
*Rpl32 Reverse:* gcc ctt gtt cga tcc gta

### Western Blotting

Whole-cell lysates were prepared from *Drosophila* heads. For each sample, 10 heads were collected on dry ice and stored at -80 until processing. Tissue was resuspended in 200 µL ice-cold RIPA buffer (150 mM NaCl, 1% Triton X-100, 0.1% SDS, 0.5% sodium deoxycholate, 50 mM Tris [pH 8.0]) supplemented with protease /phosphatase inhibitor cocktail [Pierce Protease and Phosphatase Inhibitor Mini Tablets, Thermo Scientific cat:A32959]. Samples were homogenized with a pestle for 10 s, vortexed for 10 s, and incubated on ice for 10 min. Lysates were cleared by centrifugation at 16,000 × *g* for 10 min at 4 °C, and the supernatant was collected. Protein concentration was determined using a BCA assay [Pierce BCA Protein Assay Kit, Thermo Fisher], and samples were normalized to equal concentrations.

Lysates were mixed 4:1 with 4x Laemmli sample buffer containing 200 mM DTT and denatured at 70 °C for 10 min. 20 µg of protein per lane was separated by SDS-PAGE on 8 or 10% gels in Tris/glycine/SDS running buffer at 125 V for 60 min. Proteins were transferred to 0.2 µm PVDF membranes by wet transfer or dry transfer (Bio-Rad Turbo Blot) in cold Tris/glycine buffer containing 20% (v/v) methanol at 70 V for 1 h at 4 °C. PVDF membranes were activated in 100% methanol prior to transfer. Transfer efficiency was verified by No-Stain Protein Labeling Reagent (Invitrogen cat:A44449).

Membranes were incubated with primary antibodies overnight at 4 °C. Membranes were washed three times for 15 min in TBS-T, incubated with HRP-conjugated secondary antibodies (Mouse, Rabbit Cell Signaling 1:10000) in 5% nonfat milk for 1– h at room temperature, and washed again three times for 15 min in TBS-T. Signal was detected using Prometheus ProSignal Pico (Genesee Scientific) and imaged on an Invitrogen iBright. Band intensities were quantified using Bio-Rad Image Lab software and normalized to total protein.

Primary antibodies used were: anti-GRP78/BiP (rabbit 1:1000: Proteintech, catalog # 66574-1), anti-Thr-AMPylation (mouse 1:1000, BioIntron, 17G6), anti-DOPR1 (rabbit 1:000, Invitrogen, Thermo Scientific, catalog # PA5-144383).

### Brain Immunohistochemistry

Adult brain immunostaining was performed as previously described(44). Adult brains were dissected and fixed in PBS, pH 7.4 containing 3.7% formaldehyde for approximately 1 hr. Fixed brains were washed quickly 3 times followed by 3 – 20 min washes in 0.5% PBS-T, with gentle rocking at room temperature. Brains were blocked using 5% normal goat serum (NGS) in 0.5% PBS-T for at least 30 min at room temperature with gentle rocking, then incubated in primary antibody diluted in 5% NGS for three nights at 4°C. After primary antibody incubation, brains were washed 3 times in PBS-T and incubated in secondary antibody diluted 1:500 in 5% NGS for one night at 4°C. Brains were washed 3 times in PBS-T and mounted between a glass microscope slide and a #1.5 cover glass separated by a custom bridge in VECTASHIELD Antifade Mounting Medium with DAPI (Vector Laboratories). Samples were imaged on a Leica STELLARIS Confocal Microscope using a 20X air and 63X oil objective. All treatments were mounted under the same cover slip and at least two slides per experiment were imaged. Image processing was performed using ImageJ (NIH). Images are representative maximum intensity projections compiled from 1-2 μm thick sections of the z-stacks and are contrast matched. Mouse anti-calnexin99a (DSHB 1:20) and rabbit anti-Dop1R1 (Invitrogen PA5-144383, 1:30) were used for primary antibody staining, and Alexa Fluor 488 and 633 were used for secondary antibody staining (Life Technologies 1:500).

### *C. elegans* locomotor tracking

*C. elegans* strains were maintained at 20 °C on standard nematode growth medium (NGM) plates seeded with *E. coli* OP50 and were kept well fed for at least two generations before use. Wild-type N2 and *fic-1* null [*fic-1(n5823)*] animals were each crossed into strain CZ10175 [*zdIs5*(*mec-4p::GFP*)]. Worm populations were age-synchronized by alkaline hypochlorite treatment as previously described(45). Mixed-stage animals were rinsed off plates with M9 buffer into 1.7 mL microcentrifuge tubes, allowed to settle by gravity for 3–5 min, and rinsed three to four times with fresh M9. The M9 was then replaced with 1 mL of hypochlorite solution (approximately 0.93 M NaOH and 0.70% sodium hypochlorite final), and tubes were agitated at 1,200 rpm and 20 °C on a heated shaker for 7–10 min, until no intact worm bodies remained. Released eggs were pelleted at 1,600 × g for 30 s and washed three times with 1 mL M9, followed by a final M9 wash and centrifugation at 2,900 × g for 3 min. Eggs were resuspended in approximately 100 µL M9, plated onto seeded NGM plates, and grown at 20 °C until day 1 of adulthood.

For each genotype, 20–30 day 1 adults were transferred to 35 mm NGM plates spotted with 100 µL OP50 and allowed to acclimate for 30 min. Plates were then placed in a WMicrotracker X8 SMART (Phylum Tech), and locomotor activity was recorded for 5 min in tracking mode. Output measures, e.g., average speed and distance traveled using Phylum Tech, Software Version: SMART V1.6 2023. The assay was performed in 5-6 independent biological replicates.

### Sucrose preference test

Mice were habituated to a modified homecage setup where they had access to two water bottles for 3 days prior to any experimental manipulation. Mice were then given 1 bottle of water and one bottle of either 8% sucrose or 0.8% sucralose. Preference tests started at 10AM and continued for 24 hours, with the bottle positions switched six hours into the intervention to control for mice preferring a given side of the cage. After at least one day of washout (water available in both bottles), mice then received the other tastant, with order counterbalanced across mice to control for order effects. After at least one more washout day (water available in both bottles), all mice were given one bottle of 8% sucrose and one bottle of 0.8% sucralose to assess preference between sweet tastants. The amount consumed was determined after 24 hours of access to a given pair of liquids. Preference was calculated by dividing the amount of a given liquid consumed divided by the sum of all liquid consumed.

### Statistics

For comparisons involving only one level, we used Student’s t-test to detect significant differences between two treatments or one-way ANOVA followed by Tukey’s post-hoc test after verifying normality and equality of variances. T-tests were two-tailed during initial characterization experiments, or one-tailed in future experiments where the predicted direction of change was known. For comparisons involving more than one level, we used two-way ANOVA to detect significant interactions between the levels and followed up with Tukey’s post-hoc when significance was detected (p<0.05). In cases where experimental replicates were pooled, a two-way ANOVA with blocking for experiment was performed to ensure non-significant experimental effects. For all dot and bar plots, error bars represent the SEM. Statistical tests and graphing were performed using GraphPad Prism or R Studio. Specific details of statistical analyses are presented in the figure legends, where *n* per treatment is also listed.

## Acknowledgments

We thank the members of the Pletcher and Truttmann labs for their help with this collaborative effort. Thank you to Kim Orth and Helmut Kramer for the kind sharing of resources and advice, and to the Burgess lab for their contributions. AI (Claude Opus 5.5) was used to draft and edit this manuscript, and for the generation of code to analyze FLIC and RNA-sequencing data.

## Author Contributions

Designed research: K.J.W., B.A., C.B., S.D.P., M.T.

Performed research: K.J.W., B.A., W.G., B.T., A.J.

Contributed new reagents or analytic tools: S.P.

Analyzed data: K.J.W., B.A., B.T., C.B.

Wrote the paper: K.J.W., B.A.

