## Supplementary Information for "Neuronal FICD shapes motivated behavior through modulation of dopamine receptor dynamics"

**This PDF file includes:**

Supplementary Figures 1-6
Supplementary Table 1

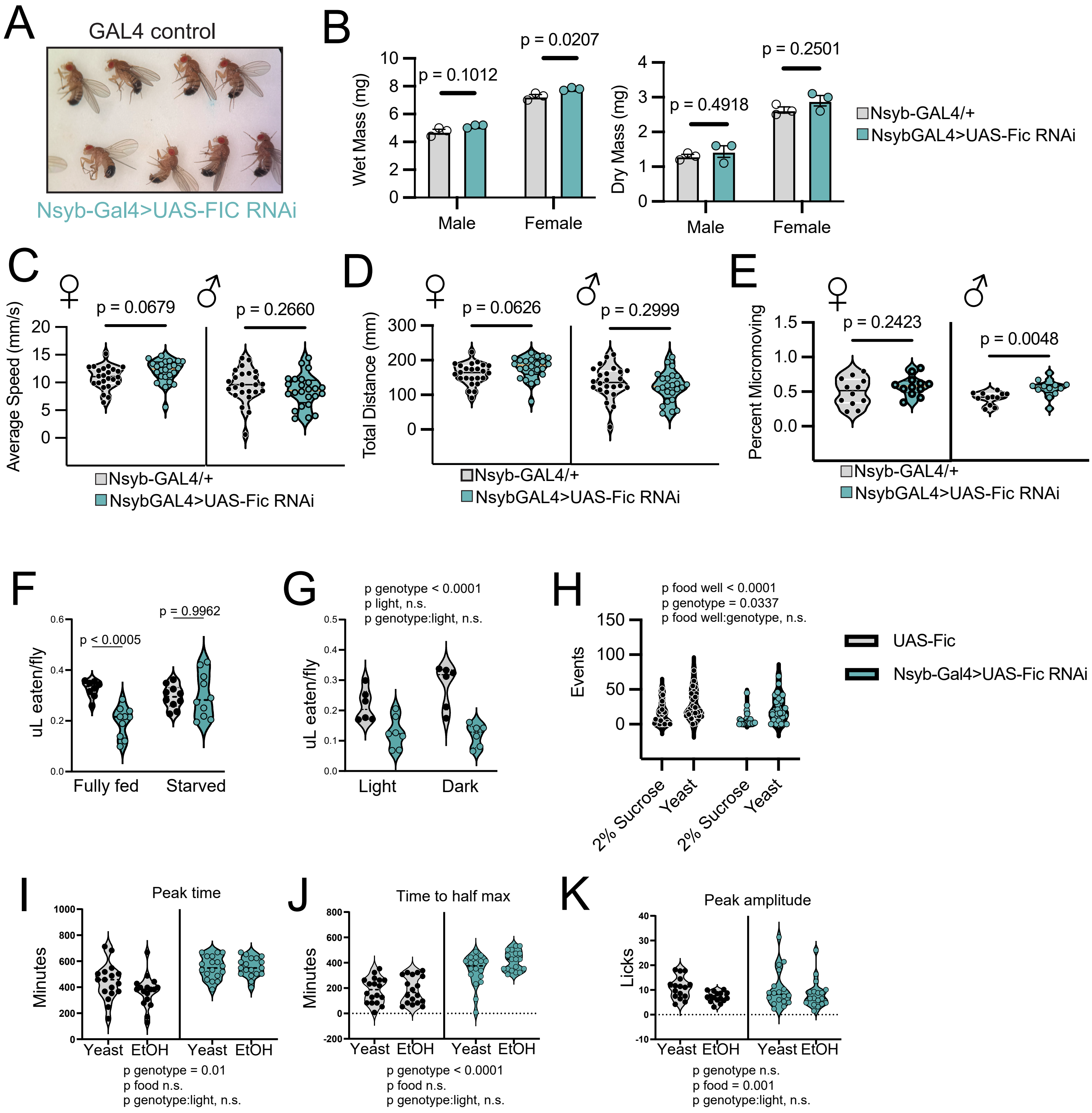


Supplementary Figure 1

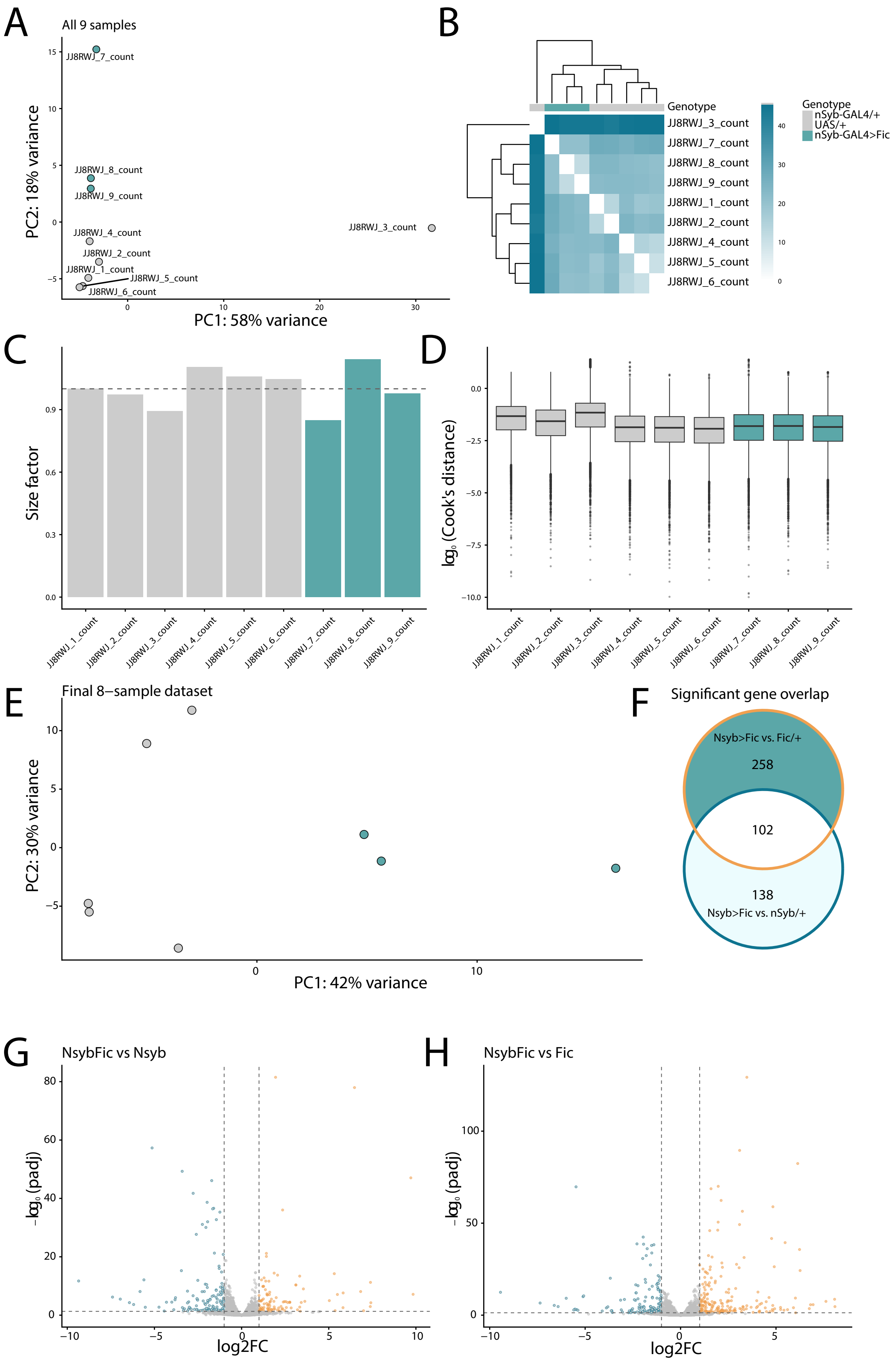


Supplementary Figure 2

A

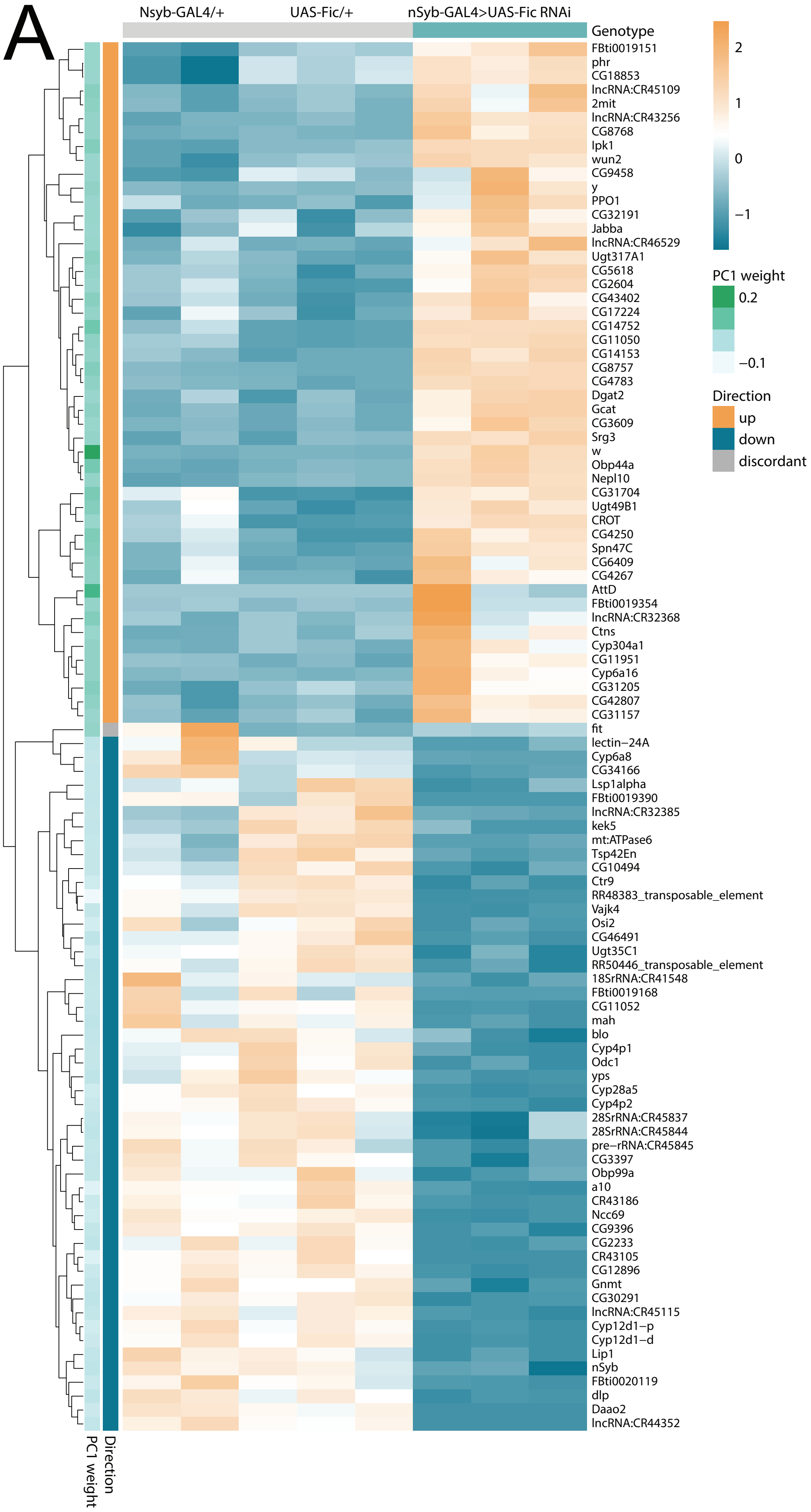


Supplementary Figure 3

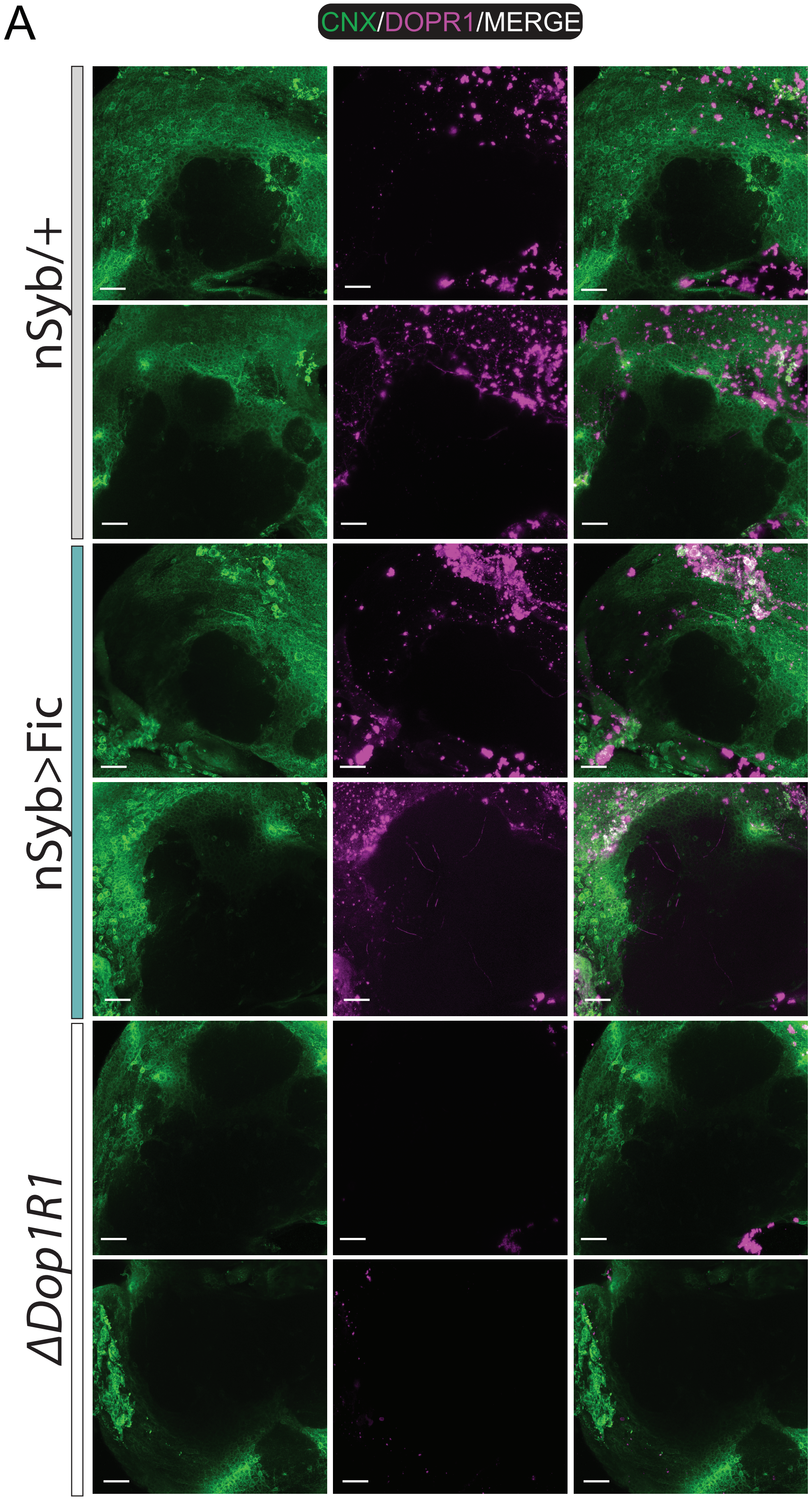


Supplementary Figure 4

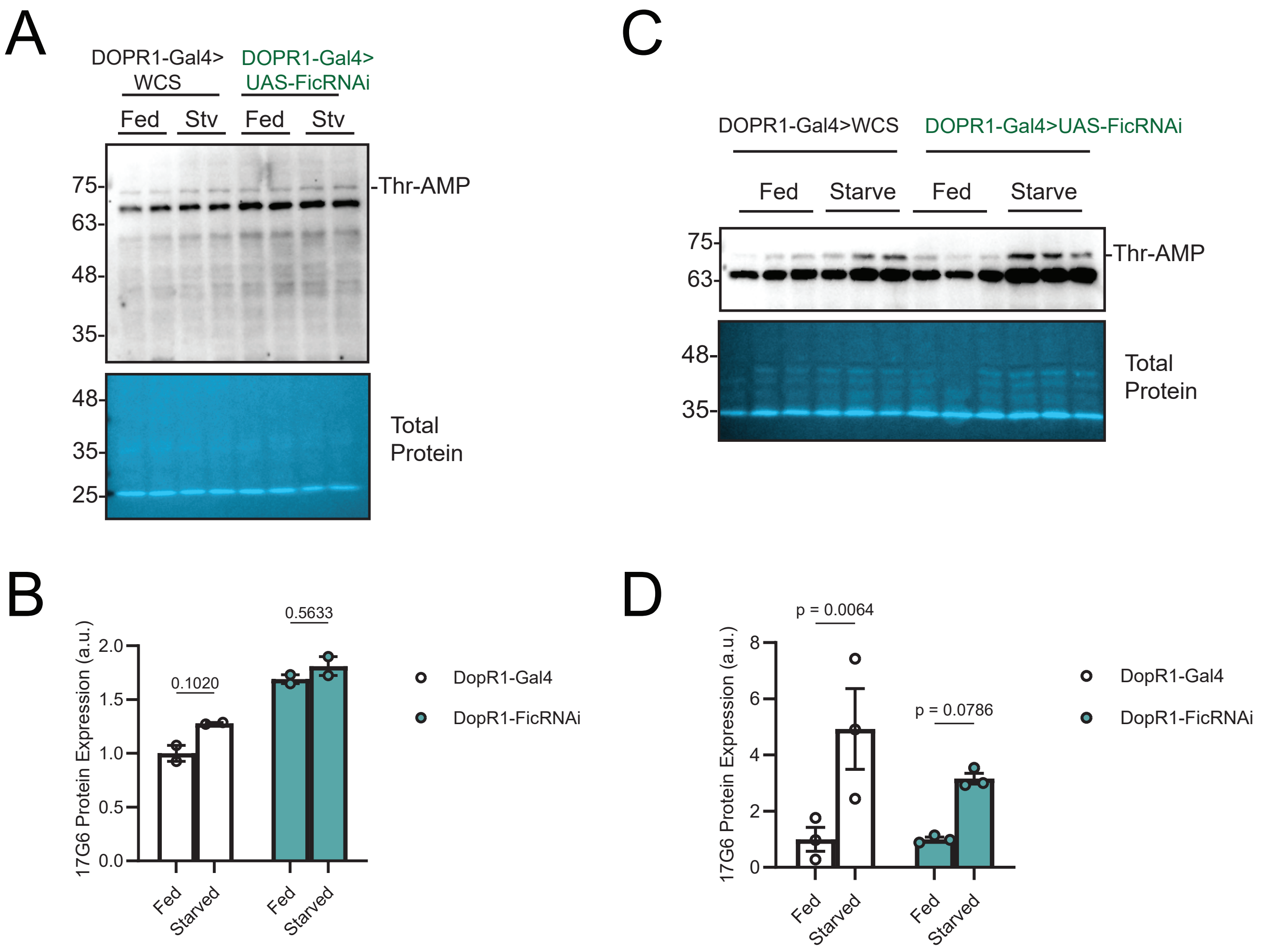


Supplementary Figure 5

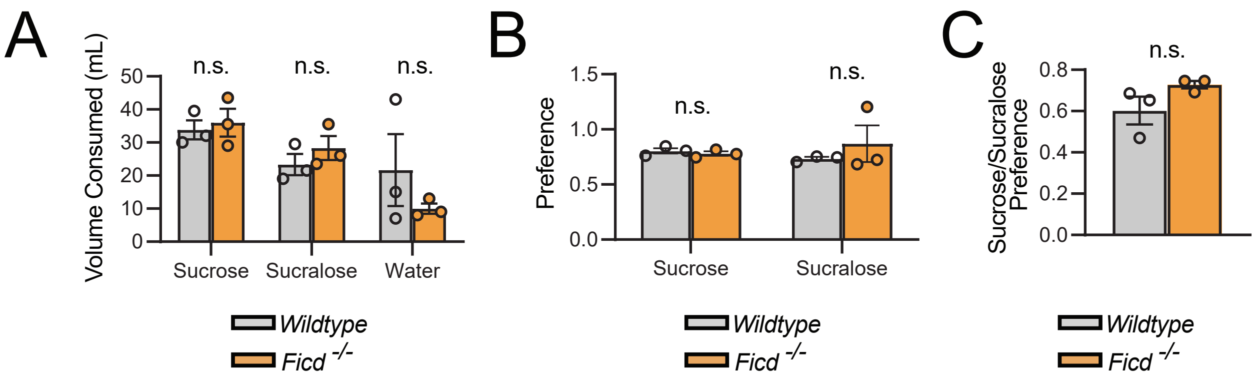


Supplementary Figure 6

### Supplementary Table 1

| Gene | FlyBase ID | log2FC (nSyb>Fic vs nSyb-GAL4/+) | padj (Nsyb>Fic vs nSyb-GAL4/+) | log2FC (nSyb>Fic vs UAS-Fic/+) | padj (nSyb>Fic vs UAS-Fic/+) | Sig. |
|---|---|---|---|---|---|---|
| *Ire1* | FBgn0261984 | -0.12 | 0.740 | -0.13 | 0.670 | |
| *PEK* | FBgn0037327 | 0.36 | 0.050 | 0.16 | 0.490 | |
| *Atf6* | FBgn0033010 | -0.11 | 0.708 | 0.00 | 0.995 | |
| *Xbp1* | FBgn0021872 | -0.19 | 0.202 | -0.11 | 0.502 | |
| *crc* | FBgn0000370 | 0.00 | 0.998 | 0.10 | 0.576 | |
| *Hsc70-3* | FBgn0001218 | -0.38 | 0.011 | -0.18 | 0.327 | * |
| *Pdi* | FBgn0286818 | -0.06 | 0.839 | -0.06 | 0.811 | |
| *Hrd3* | FBgn0028475 | 0.21 | 0.444 | 0.13 | 0.653 | |
| *sip3* | FBgn0039875 | -0.16 | 0.550 | -0.04 | 0.925 | |

*log2FC and Benjamini-Hochberg adjusted p-values (padj) are shown for nSyb>Fic relative to each control genotype (nSyb-GAL4/+ and UAS-Fic/+), from DESeq2 results. None of the nine genes reached significance in both comparisons (padj < 0.05). * indicates padj < 0.05 in the indicated comparison only.*

**Supplementary Figure 1. Reduced motivated behavior in nSyb>Fic flies is not explained by developmental, motor, sensory, or homeostatic deficits.**

**(A)** Representative image of GAL4 control and Nsyb-Gal4>UAS-Fic RNAi adults, showing grossly normal external morphology. **(B)** Wet mass (mg, left) and dry mass (mg, right) of males and females [n = 3] (unpaired two-tailed t-test). **(C)** Average speed (mm/s) from high-resolution video tracking of freely moving females and males [n = 12] (unpaired two-tailed t-test). **(D)** Total distance traveled (mm) from the same tracking (unpaired two-tailed t-test). **(E)** Percent of time spent micromoving (unpaired two-tailed t-test). **(F)** FLIC food intake (µL eaten/fly) under starved conditions; knockdown flies eat less than controls when fully fed ($p < 0.0001$) but equivalently after starvation ($p = 0.8274$), showing that maximal feeding capacity is intact (two-way ANOVA with tukey's, interaction $p = 0.0018$). **(G)** FLIC food intake (µL eaten/fly) tested in the light and dark; knockdown flies eat less than controls under both conditions (two-way ANOVA). **(H)** FLIC interaction events with 2% sucrose versus yeast food for UAS-Fic and Nsyb>Fic; both genotypes preferentially engage the yeast food (interaction n.s.), indicating intact homeostatic drive. **(I–K)** FLIC feeding-bout dynamics on yeast versus ethanol food for control and knockdown: peak time (I, minutes), time to half-maximum (J, minutes), and peak amplitude (K). All error bars represent the mean ± SEM.

**Supplementary Figure 2. Sample quality control, outlier exclusion, and differential-expression overview for the nSyb>Fic RNA-seq dataset.**

**(A)** Principal-component analysis (PCA) of variance-stabilizing-transformed (VST) counts for all nine sequenced samples (Nsyb-GAL4/+, n = 3; UAS-Fic/+, n = 3; nSyb-GAL4>UAS-Fic RNAi, n = 3). One Nsyb-GAL4/+ replicate separates from all other samples along PC1 (58% of total variance). **(B)** Hierarchical clustering heatmap of pairwise sample-to-sample Euclidean distances (VST counts), colored by genotype. **(C)** DESeq2 size factors per sample, colored by genotype. **(D)** Per-sample distribution of $\log_{10}$(Cook's distance) across all tested genes, colored by genotype. **(E)** PCA of VST counts for the final 8-sample dataset after exclusion of the outlier Nsyb-GAL4/+ replicate identified in (A), (B), and (D) (see Methods); PC1 = 42% of variance. **(F)** Venn diagram of significant genes (padj < 0.05, |log2FC| > 1) for nSyb>Fic vs. Nsyb-GAL4/+ (138 genes) and nSyb>Fic vs. UAS-Fic/+ (258 genes), with 102 genes significant in both; 97 genes remain after filtering to those changing in the same direction. **(G, H)** Volcano plots of differential expression for nSyb>Fic vs. Nsyb-GAL4/+ (G) and nSyb>Fic vs. UAS-Fic/+ (H); genes colored by significance and direction (padj < 0.05, |log2FC| > 1).

**Supplementary Figure 3. Differentially expressed genes in nSyb>Fic heads.**
**(A)** Heatmap of differentially expressed genes (rows, z-scored expression) across the three genotypes (columns: Nsyb-GAL4/+, UAS-Fic/+, nSyb-GAL4>UAS-Fic RNAi). Genes shown are those significant in the same direction relative to both controls (padj < 0.05, |log2FC| > 1) together with the top positive- and negative-PC1-loading genes distinguishing nSyb>Fic from controls. Row annotations indicate PC1 loading weight (green scale) and direction of change (up, orange; down, blue; discordant, grey).

**Supplementary Figure 4. Dysregulated dFic mislocalizes Dop1R1**
**(A)** Immunofluorescence of UAS-Fic RNAi/+ (control) and nSyb-GAL4>UAS-Fic RNAi brains stained for calnexin (CNX, green), Dop1R1 (DOPR1, red), and DAPI (blue). Images are maximum intensity projections of the lateral protocerebrum.

**Supplementary Figure 5. The starvation-induced rise in BiP AMPylation is blunted in the heads, but not bodies, of Dop1R1>Fic flies.**
**(A)** Western blot of head lysates probed for AMPylation (17G6), with total protein as loading control. Molecular-weight markers (kDa) at left. **(B)** Quantification of 17G6 signal (a.u.) for (A), fed versus starved, in control (DopR1-Gal4) and knockdown (DopR1-FicRNAi) (two-way ANOVA). **(C)** Western blot of body lysates, as in (A). **(D)** Quantification of 17G6 signal (a.u.) (two-way ANOVA). All error bars represent the mean ± SEM.

**Supplementary Figure 6. Female FICD$^{-}$/$^{-}$ mice show no change in sucrose consumption or preference.**

**(**A) Fluid consumption (mL) by wild-type and global Ficd$^{-}$/$^{-}$ female mice in a two-bottle test: sucrose, sucralose, and water [N = 3] (unpaired two-tailed t-test, all comparisons n.s.). **(B)** Preference (fraction of intake from the tastant bottle) for sucrose and sucralose (unpaired two-tailed t-test, both n.s.). **(C)** Sucrose/sucralose preference ratio (unpaired two-tailed t-test, n.s.). All error bars represent the mean ± SEM.
